# Protein Response to ACL Injury in Humans Show Early Cartilage Remodeling and Sex-Specific Effects

**DOI:** 10.64898/2026.05.12.724692

**Authors:** Paula A. Hernandez, Constance R. Chu, Chun-Yuan Huang, Chao Xing, Meenakshi V. Venkatachalam, J. Lee Pace, Steven B. Singleton

## Abstract

**Objective:** Anterior cruciate ligament (ACL) tears increase the risk for developing knee osteoarthritis (OA). Females have greater risk for both. However, studies defining sex-specific protein responses in human cartilage after ACL injury are lacking. We hypothesize that articular cartilage’s response to an injurious environment differs depending on sex.

**Methods:** We compared the proteomic profiles of *normal* cartilage with *injured* cartilage harvested from the intercondylar area during ACL surgery. Sex-specific injury effects were estimated through contrasts between Injured Male and Normal Male and between Injured Female and Normal Female. Over-Representation Analysis (ORA) was done using gene ontology (GO) and the Kyoto Encyclopedia of Genes and Genomes (KEGG) database. Extracellular matrix (ECM) proteins were further analyzed using the Matrisome AnalyzeR.

**Results:** From the 2,188 proteins identified, males and females shared 1,121 upregulated and 23 downregulated proteins in injured compared to normal cartilage. Analysis of ECM proteins and enriched pathways revealed mostly similar male and female responses to an injurious environment, with evidence of early cartilage remodeling in both sexes. Nevertheless, more than 240 proteins were affected specifically by sex, and significant sex differences were found in inflammation, ECM-related, and metabolic pathways. Males were enriched mostly in “*ECM-receptor interaction*”, while females were enriched in “*Citrate cycle (TCA cycle)*”, “*Fatty acid degradation*”, and “*Fatty acid metabolism*” pathways.

**Conclusion:** Articular cartilage shows signs of remodeling soon after ACL injury, even when only exposed to an injurious environment rather than being physically impacted. Sex-specific responses were observed in inflammation, metabolic pathways, and ECM synthesis.

## INTRODUCTION

Sport-related knee injuries, such as anterior cruciate ligament (ACL) tears, are becoming more frequent with the steady increase in sports participation and sports specialization at a young age^1, 2^. These injuries confer a higher risk for developing posttraumatic osteoarthritis (PTOA) later in life^3, 4^, with PTOA developing in up to 40% of cases within 10 years after ACL tear and/or ACL surgery^5^. Moreover, there exists a disparity between the sexes, with females having increased risks for both sustaining ACL tears^6–9^ and developing knee OA^10, 11^. The cause of the differing risks and rate of injury profiles between males and females seems multifactorial and includes anatomical, biomechanical, and biological factors ^12–15^.

Comprehension of the response of knee articular cartilage to injury is evolving^16–20^ leading to more questions about changes to the extracellular matrix (ECM) and modulation of molecular pathways. Furthermore, as the whole joint is exposed to a post-injury environment, there is no complete understanding of whether normal or intact looking cartilage specimens from an injured joint can truly be considered normal. This is a relevant question, as limitations in obtaining normal young human cartilage for research to be used as control have inevitably pushed the field to use either intact looking cartilage from OA samples^21^ or non-load bearing biopsies obtained at the time of reconstructive surgery (non-OA)^22, 23^. Although these studies have allowed the discovery of some of the early degenerative changes occurring in cartilage, it has been observed that normal looking cartilage from early onset hip OA already had OA-like cellular changes, even though the macrostructure of the tissue still appeared intact^24^, indicating that the joint environment can influence intact nearby tissues^25, 26^.

Moreover, age and sex are factors generally not considered in the analysis of many studies. Indeed, whether there are sex-specific changes in articular cartilage post knee injury in humans is understudied. Unveiling the events affecting the ECM in human cartilage after an ACL tear and before the onset of PTOA, specially from the sex-specific perspective can be beneficial not only for the advancement in women’s health, but also for developing better strategies to manage cartilage repair in young athletes regardless of biological sex.

Animal models of PTOA have provided us with some improved understanding of what occurs at the ECM and molecular levels in OA development and are crucial for drug discovery due to its translatability to human health. In particular, rodent models of PTOA have allowed for longitudinal studies with more homogenous outcomes compared to human. Research using these small animal models has revealed some of the main molecular pathways triggered at the cartilage level, mostly related to inflammation and pain^27, 28^. However, rodent models for PTOA have failed to recapitulate the sex differences observed in humans^28, 29^. Male mice develop faster cartilage degeneration compared to female mice, which has prompted researchers to exclude female mice from several studies. Others focusing on females have shown that sex hormones, such as estrogen, are a key component in the protection of cartilage from degeneration, as ovariectomized mice develop faster PTOA changes than intact females^30, 31^.

Therefore, the aim of this study was to unveil whether there are early changes in human articular cartilage at the ECM and molecular levels after an ACL injury and whether these events differ depending on biological sex. Our goal is to understand changes occurring after injury, but before any development of PTOA. We hypothesize that early ECM remodeling can be detected in cartilage after ACL injury and that these changes have a sex-specific component.

## METHODS

### Study Approval

Study approval to collect surgical specimens was obtained from the Stanford University Institutional Review Board (IRB 27369). Written consent was obtained from patients having ACL reconstruction (ACLR). All samples were obtained by the same surgeon, snap frozen, and stored in −80°C till use. All the de-identified tissue was transferred simultaneously to UT Southwestern under a material transfer agreement. Use of de-identified discarded surgical samples at UT Southwestern was exempt from Institutional Review Board approval. Normal cartilage from healthy post-mortem donors was provided by AlloSource from research consenting donors, after passing serology screening (6-10 days from donor’s death to receiving the tissue in the laboratory) and in full compliance with all applicable rules and regulations. Use of de-identified post-mortem samples at UT Southwestern was exempt from Institutional Review Board approval.

### Study Design and Tissue Source

We compared the proteomic profiles of *normal* cartilage (5 males, 5 females, 37-39 years old) with *injured* cartilage (5 females, 25-39 years old, BMI 23.47 ± 2.64 kg/m^2^ and 5 males, 22-39 years old, BMI 25.04 ± 3.86 kg/m^2^). Injured cartilage samples were small full thickness cartilage biopsies harvested from the lateral notch in the intercondylar area during ACLR. Details of the samples used in this study are summarized in Table 1 and Supplementary Table 1. Normal cartilage were full-thickness shavings from the intercondylar and condylar areas from healthy donors (without musculoskeletal diseases). To limit the aging effect and the hormonal decrease after menopause, all samples used in our study come from individuals aged 22 to 39 years old.

**Table 1:** Summary of demographic data of all injured samples used in this study.

| Sample Type | Sex | Age | Sample number | BMI kg/m<sup>2</sup> | Months since injury | Meniscus tear? | MCL/LCL tear? | Articular defect? | First time ACL tear? | Injury mechanism? |
| --- | --- | --- | --- | --- | --- | --- | --- | --- | --- | --- |
| Injury | M | 22 | 1 | 30.74 | 6 | LMT | no | no | yes | non-contact |
| Injury | M | 26 | 2 | 25.77 | 1.5 | LMT | no | no | yes | non-contact |
| Injury | M | 29 | 3 | 20.12 | 2 | LMT | MCL | no | yes | non-contact |
| Injury | M | 29 | 4 | 23.51 | 3.5 | no | no | no | yes | non-contact |
| Injury | M | 39 | 5 | 25.08 | 2 | LMT, MMT | no | MCF | yes | non-contact |
| Injury | F | 25 | 6 | 24.65 | 1.5 | LMT, MMT | no | no | yes | non-contact |
| Injury | F | 26 | 7 | 24.33 | 1.5 | LMT, MMT | MCL | no | yes* | non-contact |
| Injury | F | 27 | 8 | 22.06 | 0.5 | no | no | no | yes | non-contact |
| Injury | F | 31 | 9 | 19.71 | 3 | LMT | no | no | yes | non-contact |
| Injury | F | 39 | 10 | 26.58 | 2 | no | MCL | no | yes | contact |

LMT = lateral meniscus tear, MMT = medial meniscus tear, MCL = medial collateral ligament, MFC = medial femoral condyle; \*prior contralateral ACLR

### Proteomics Assay and Analysis

All samples were processed at the same time. Specimens were snap frozen, pulverized in liquid nitrogen, and resuspended in 5% sodium dodecyl sulfate (SDS) with 50 mM of triethylammonium bicarbonate (TEAB) in a ratio of 500 µl per 100 mg of powder. A pooled standard was made using equal volumes from each sample to be used downstream in data processing for batch-effect correction. labeled Equal amounts of peptide per sample based on a Nanodrop A205 reading of peptide concentration were labeled using Tandem Mass Tag (TMT). The two sets of TMT data were normalized with the batch effects accounted for using internal reference scaling (IRS)^32^. The false discovery rate cutoff was 1% for all peptides. Proteomic analysis was done at the UT Southwestern Proteomics Core Facility as previously described^33^.All analyses were conducted in R with the limma, DEqMS, ggplot2, pheatmap, and VennDiagram packages. Prior to statistical comparisons, proteins were filtered to require values in at least 50% of all the runs. The data was log_2_-transformed and imputed replacing values from a down-shifted normal distribution, using default settings within Perseus 2.0.9.0. Multidimensional scaling (Euclidean distance) was used to visualize overall sample clustering, with groupings by sex and injury status. Sex-specific injury effects were estimated through contrasts between Injured Male and Normal Male and between Injured Female and Normal Female. Differential abundance was assessed using linear modeling framework (limma package) that incorporated cell means for the four experimental groups (Normal Male, Normal Female, Injured Male, Injured Female) and included batch as a covariate^34^. Variance estimates were refined using DEqMS package after limma, which models protein-level standard errors based on peptide-spectrum match counts to improve accuracy in differential analysis^35^. P-values were adjusted for multiple testing using the Benjamini–Hochberg method to control the false discovery rate (FDR < 0.05)^36^.

### Enriched Pathways Analysis

Differentially expressed proteins between normal and injured cartilage per sex were considered for Over-Representation Analysis (ORA) when log_2_ fold change was <−1 or >1 and with adjusted p-value <0.05.

Enrichment analysis was done using gene ontology (GO) and the Kyoto Encyclopedia of Genes and Genomes (KEGG) database with ShinyGO V0.85^37^. Extracellular matrix (ECM) proteins within the differentially expressed proteins per each sex were identified and further analyzed using the Matrisome AnalyzeR^38^. A second enrichment analysis was done considering only those differentially expressed proteins that were not shared between male and female cartilage (sex-specific analysis). These targets had a log_2_ fold change of <−1 or >1, and an adjusted p value <0.05.

### Data Availability

All data is included in this manuscript and will be uploaded to MassIVE after publication.

## RESULTS

### Shared responses between males and females

The time post-injury of the samples showed no significant difference between males and females (Fig. 1A). A total of 2,188 proteins were identified on the proteomic analysis. From those, 1,660 were either up or down regulated in injured cartilage compared to normal cartilage. Principal component analysis indicates that samples separate mostly by condition (injured vs normal) rather than by biological sex suggesting that male and female cartilage share similar responses to injury (Fig. 1B-C). This was confirmed by clustering analysis of differentially expressed proteins by sex. The abundance trend of two major clusters of proteins appears similar between normal males and females shifting to opposite expression levels in injured tissue. This shift to opposite expression levels occurred similarly in both males and females (Fig. 1D). Indeed, from the 1,660 proteins affected, male and female shared 1,121 upregulated and 23 downregulated proteins in injured cartilage compared to normal cartilage (Fig. 1E). However, there were 252 and 19 proteins that were upregulated and downregulated respectively only in males, and 242 and 3 proteins that were upregulated and downregulated respectively only in females (Fig. 1E). A heatmap of the proteins most significantly altered with injury as well as a volcano plot are shown in Figure 1F-G for male cartilage and in Figure 1H-I for female cartilage. Among the downregulated genes with injury in both sexes were structural components of the ECM.

**Figure 1:**
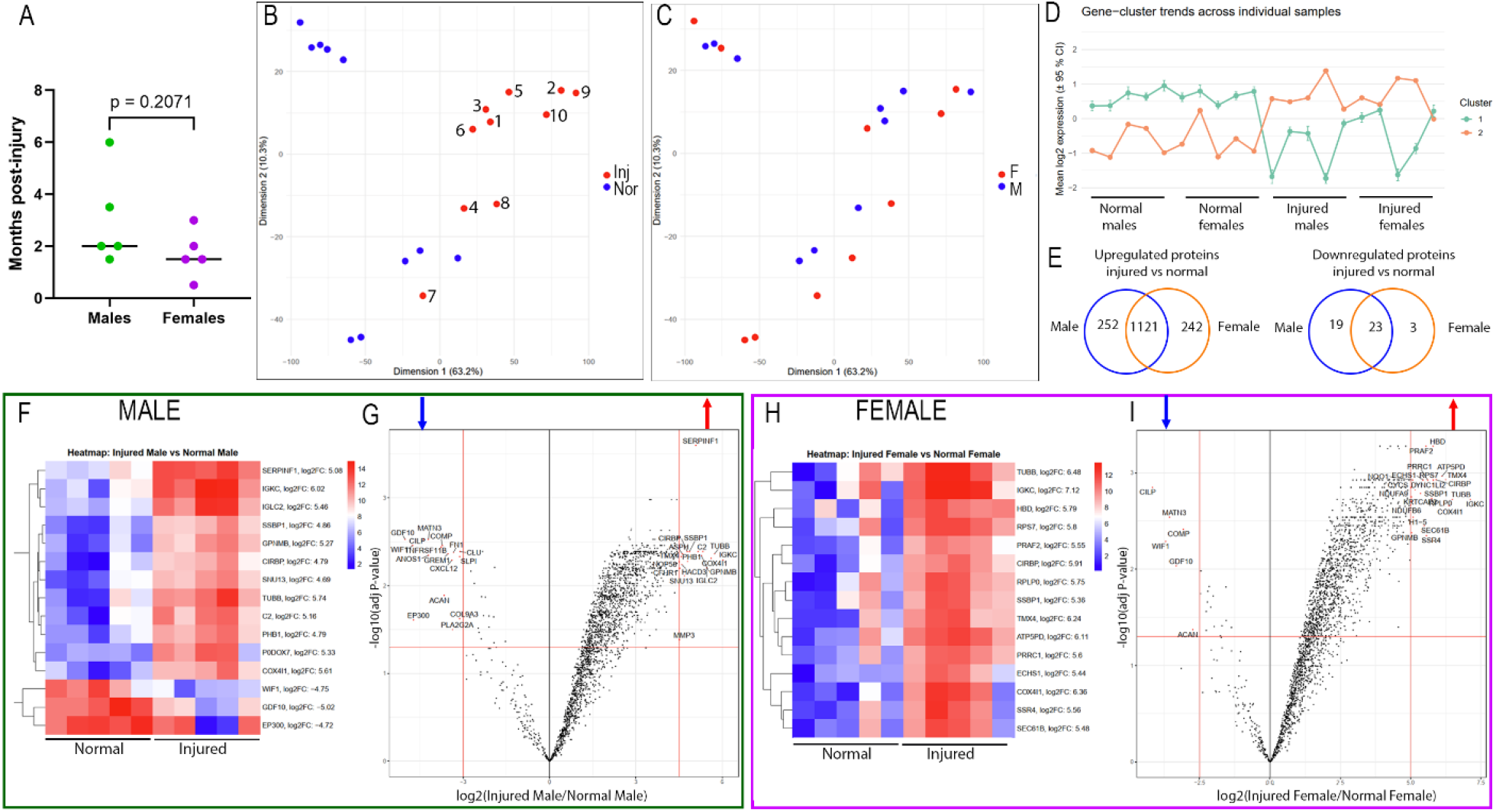
Most of the differences were detected between normal and injured cartilage. (A) Comparison of the time post-injury in male and female samples. Data is shown as individual data points with median value. P value was calculated with a two-tailed unpaired Welch’s t-test. (B) Principal component analysis shows the separation by condition and (C) by sex. Sample number of injured group from Table 1 are shown in B. (D) Cluster analysis of differentially expressed proteins further shows similarities between sexes and separation by condition. (E) Venn diagram showing shared upregulated and downregulated proteins. In both sexes there were more than 240 sex-specific proteins upregulated and less than 20 downregulated. (F-G) Heat map and volcano plot showing the most upregulated (red) and downregulated (blue) proteins with FDR <0.05 in injured compared to normal cartilage in males. (H-I) Heat map and volcano plot showing up- and downregulated proteins in injured compared to normal cartilage in females. Proteins with FDR <0.05 are shown above the horizontal red line. For clarity, only 15 proteins with the highest fold change are labeled in the volcano plot and the same representative proteins were plotted in the heat map.

### Changes in ECM-related proteins in males and females

From the pool of up- or downregulated ones in injured cartilage, we utilized the Matrisome AnalyzeR to identify which proteins are structural or ECM-related and categorize them in two groups *Core matrisome* and *Matrisome-associated*. Within the *Core matrisome* group, proteins can be further divided into proteoglycans, ECM glycoproteins, and collagens. *Matrisome-associated* proteins can be classified into ECM-affiliated proteins and secreted factors. Male and female cartilage had a similar distribution of up- or downregulated proteins in injury that belong to any of these categories (Fig 2A). However, males had a higher number of combined *Core matrisome* plus *Matrisome-associated* proteins affected with injury, with a total of 100 versus 70 for females. Additionally, slightly more ECM glycoproteins and ECM regulators were found in injured male cartilage compared to injured female cartilage. Proteins uniquely regulated per sex were 44 for males and 14 for females, and 56 in common (Supplementary Tables 2 and 3).

**Figure 2:**
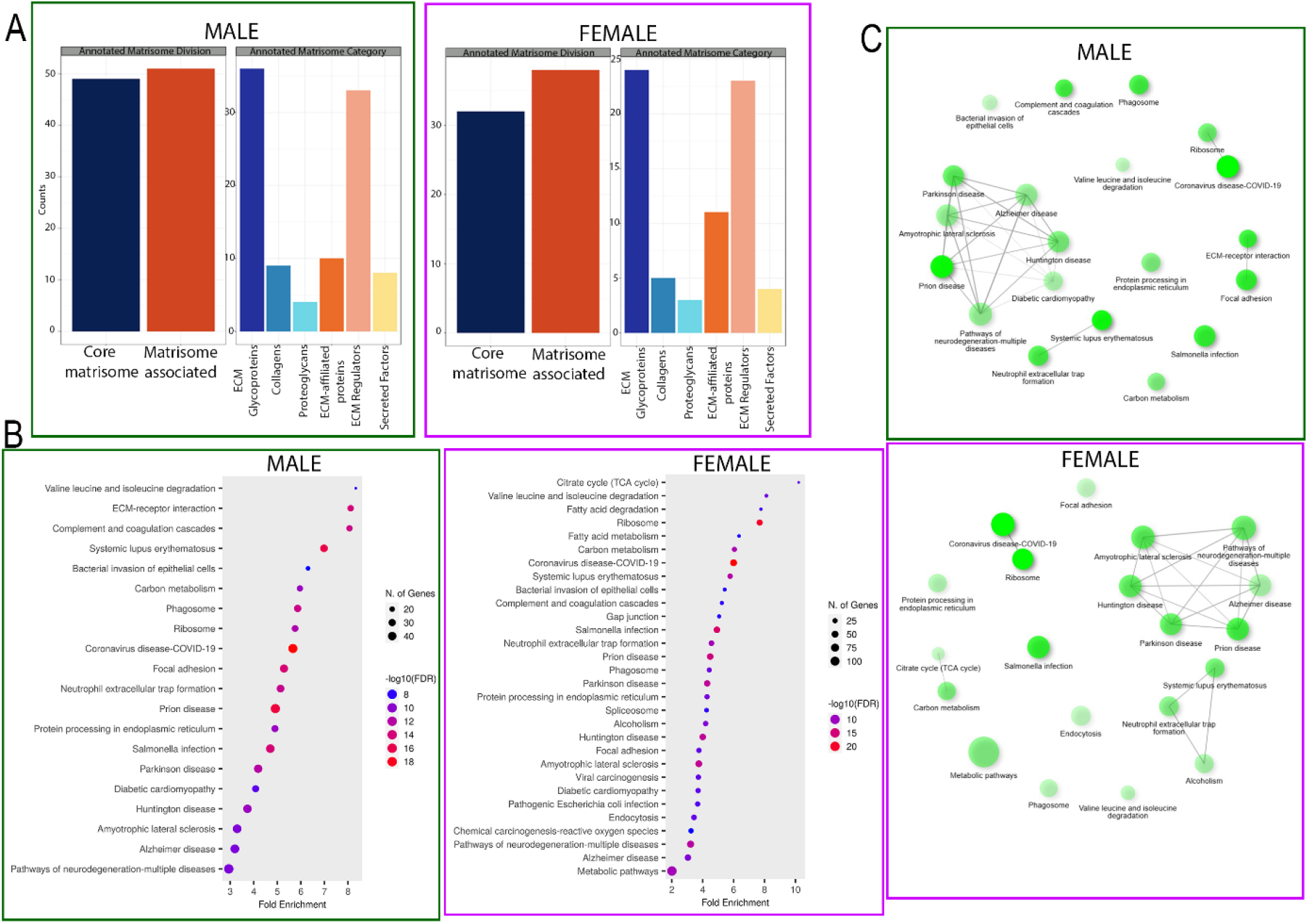
Similarities in affected ECM-related targets in both sexes and differences in enriched pathways. (A) Analysis of all up- and downregulated proteins in the Matrisome database revealed a similar distribution of ECM-related proteins in males compared to females. However, a greater number of ECM-related proteins (Core Matrisome + Matrisome Associated) were affected in male injured cartilage compared to female injured cartilage (100 vs 70). (B) GO pathway analysis of male and female injured cartilage compared to normal cartilage which are further highlighted in the network pathway analysis (C).

Among those upregulated in injured cartilage in both sexes, we found collagens type I, IV, V, VI, transforming growth factor beta-1 (TGFB1), the matrix proteases MMP3, 14, 19, and ADAM10, and S100A10, which participates in the immune response. On the other hand, ACAN, COMP, MATN3, CILP, FN-1, GDF10, the serine protease HTRA1, the chemokine CXCL12, and the WNT inhibitory factor 1 (WIFI1) were among the downregulated proteins in both male and female injured cartilage (Table 2). Although not classified as ECM related proteins in the MatrisomeDB^2.0^ database, we found that the receptors DDR2 (Discoidin domain receptor 2) and CD44 were also upregulated with injury in both sexes. Other interesting, upregulated targets in both sexes are the molecular chaperones Heat Shock Protein 90 Alpha (HSP90AA1) and Beta (HSP90AB1), and the co-chaperone Prostaglandin E Synthase 3 (PTGES3). HSP90AA1 is inducible and HSP90AB1 is constitutively activated; both act in response to several types of cellular stress including oxidative stress^39^.

**Table 2:**
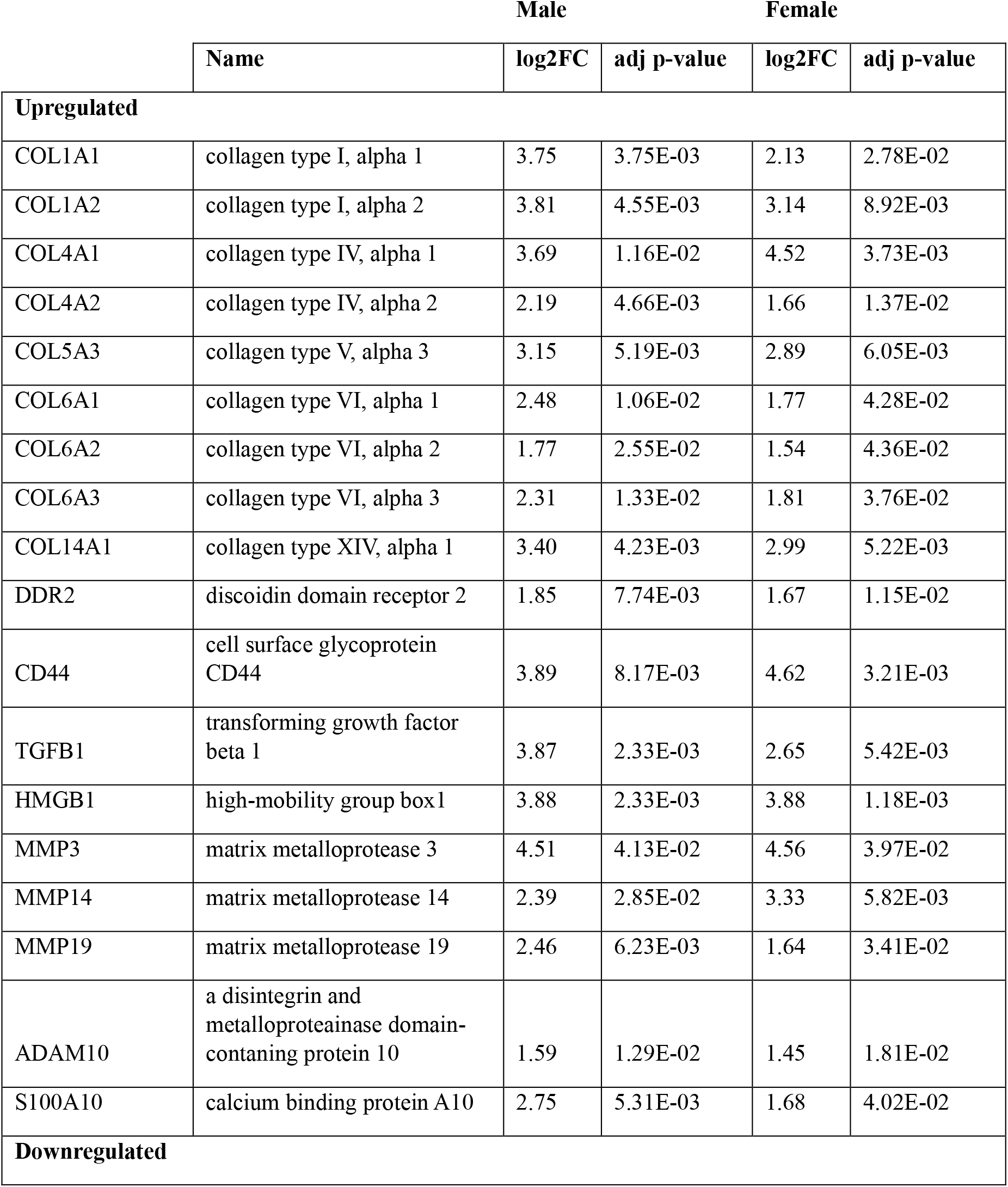
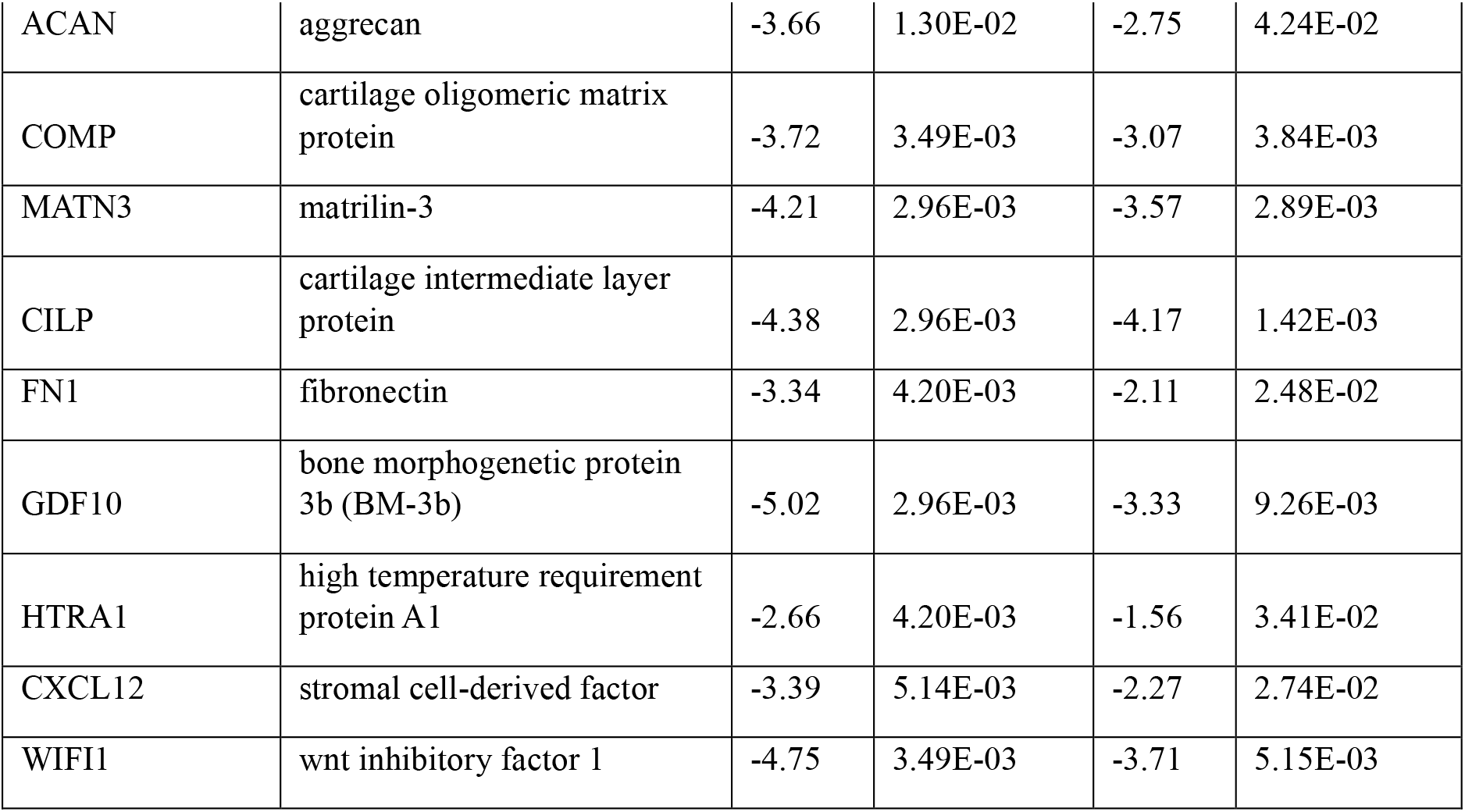
List of relevant common ECM-related proteins altered in injured cartilage in both sexes with their sex-specific log_2_ fold change (FC) and FDR.

### Over-representation analysis reveals similarities and subtle differences between sexes

KEGG pathway analysis of all differentially expressed proteins in injured samples further revealed similarities between sexes. Both males and females shared enrichment in pathways related to protein metabolism such as “*Protein processing in endoplasmic reticulum”*, “*Ribosome”*, “*Phagosome”*, “*Carbon metabolism”*, and “*Valine leucine and isoleucine degradation*”, pathways involved in inflammation such as *“Complement and coagulation cascades” and “Neutrophil extracellular trap formation”*, and pathways related to cell adhesion such as *“Focal adhesions”*. (Fig. 2B).

Nevertheless, significant sex differences were found in some pathways. While males were enriched in “*ECM-receptor interaction*”, females were enriched in energy consumption pathways such as *“Metabolic pathways”*, “*Fatty acid metabolism*”, “*Fatty acid degradation*”, and “*Citrate (TCA cycle)”*, and other cellular functions such as “*Gap junction*”, “*Spliceosome*”, and “*Endocytosis*” (Fig. 2A). Network analysis revealed that in males, there is a connection between *“ECM-receptor interaction”* and *“Focal adhesion”*. This was absent in the female analysis, and instead there was connection between *“Citrate cycle (TCA cycle)”* and *“Carbon metabolism”* that was not observed in males (Fig. 2C; Supplementary Tables 4, 5). To further explore these sex differences, we performed an enrichment analysis of those proteins, which were up- or downregulated in injured samples but were not in common in males and female samples. We found that those proteins uniquely affected in male cartilage with injury were enriched in pathways related to inflammation and interaction between cells and ECM. On the other hand, those targets affected specifically in female cartilage were related to metabolism (Fig. 3A; Supplementary Tables 6, 7).

**Figure 3:**
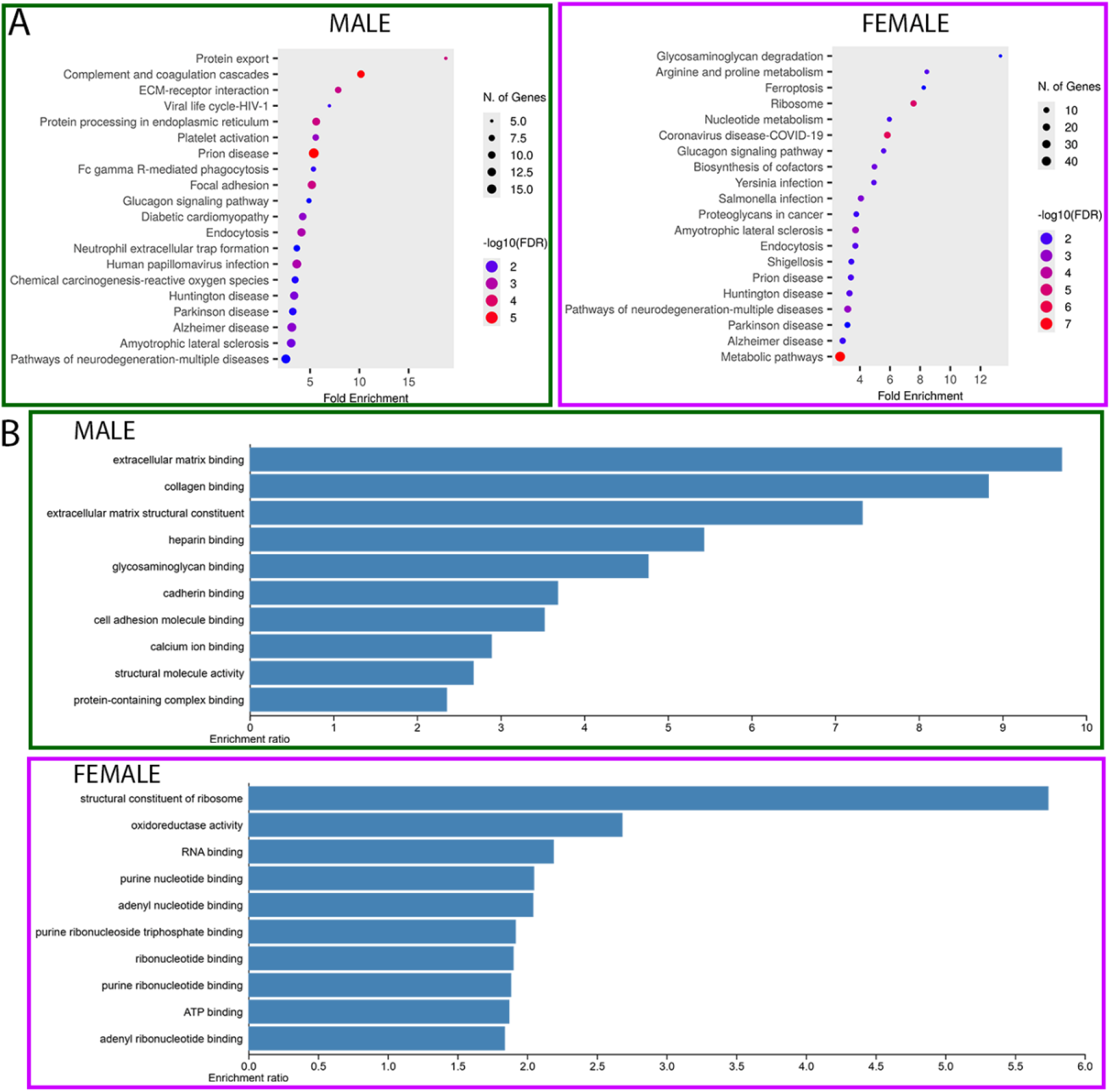
Sex-specific enriched pathways and molecular functions. (A) KEGG pathways analysis of only sex-specific up- and downregulated proteins showing sex-specific enriched pathways. (B) GO analysis of Molecular functions of only sex-specific up- and downregulated proteins. All molecular functions shown here have an FDR < 0.05.

Molecular functions of these sex-specific proteins further show that injured male cartilage is enriched in ECM-related and cell-adhesion related functions, as injured female cartilage appears enriched in metabolic-related functions (Fig. 3B). More investigation is needed to reveal whether the net balance of these functions is anabolic or catabolic.

### Sex-specific proteins

Among the group of proteins up- or downregulated with injury but unique per each sex, we found several targets relevant to cartilage biology. Among those, ECM-related and male-specific upregulated proteins included Metallopeptidase domain 9 (ADAM9), Collagen type V alpha 1 (COL5A1), Fibulin 1 (FBLN1), Matrix metalloprotease 2 (MMP2), Nidogen 2 (NID2), Integrin binding sialoprotein (IBSP), several Serpins (SERPINA1, SERPINA3, SERPIND1, SERPING1), Vitronectin (VTN), and Cartilage acidic protein 1 (CRTAC1). Male-specific downregulated proteins included Collagen IX alpha 3 (COL9A3), Cartilage intermediate layer protein 2 (CILP2), and Tissue inhibitor of metallopeptidase 3 (TIMP3). Among the ECM-related female-specific targets, Cartilage associated protein (CRTAP), Dermatan sulfate epimerase (DSE), Syndecan 2 (SDC2), and Slit guidance ligand 3 (SLIT3) were upregulated, while only Latent transforming growth factor beta binding protein 1 (LTBP1) was downregulated. (Fig. 4).

**Figure 4:**
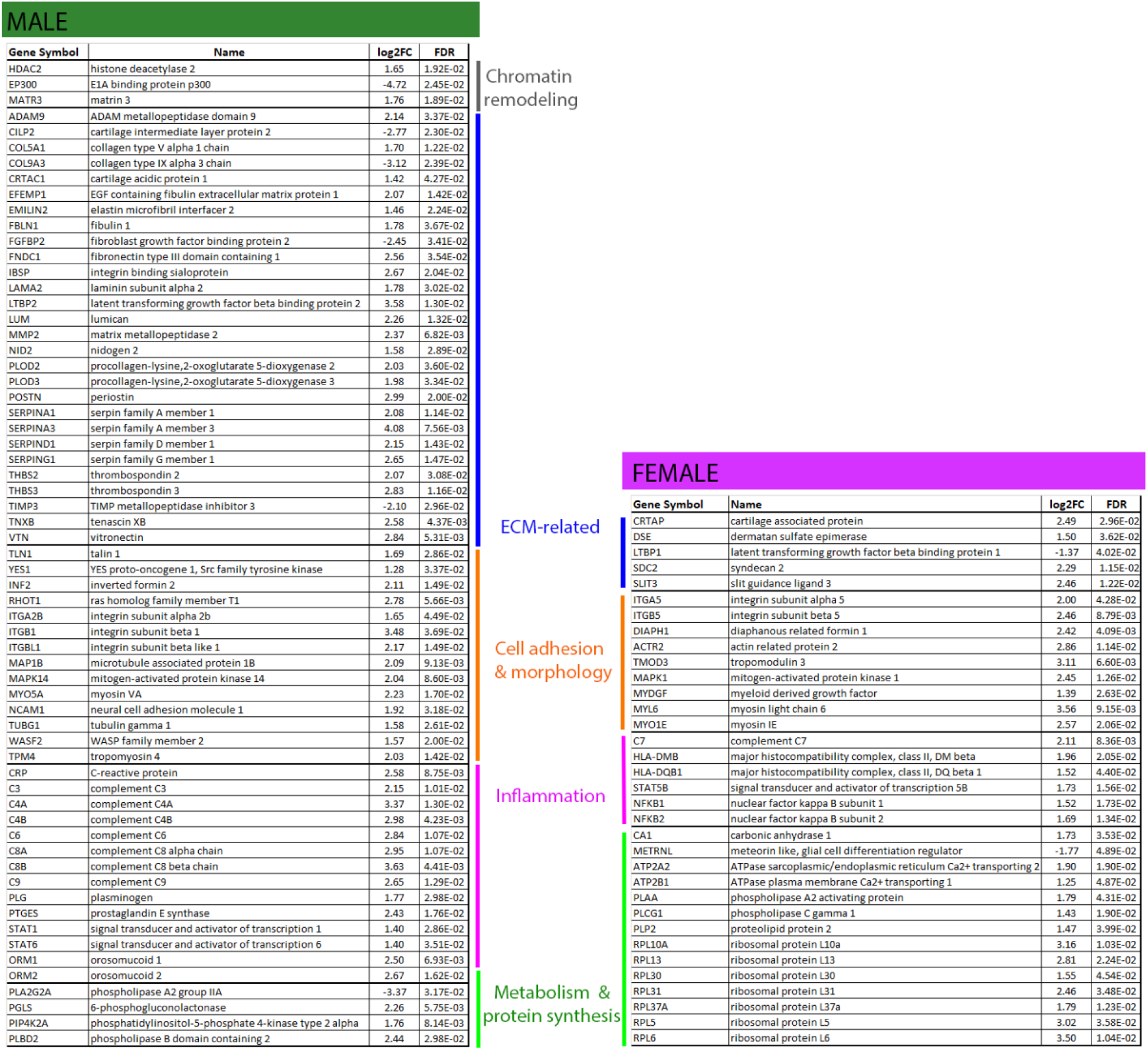
Sex-specific up- and downregulated proteins relevant for several cellular functions. List of sex specific proteins up- or downregulated in injured compared to normal cartilage. Proteins were manually classified into functions ECM-related, cell adhesion & morphology, inflammation, and metabolism & protein synthesis for comparison.

Some proteins involved in the inflammatory and immune response upregulated in males were several members of the complement system such as C-reactive protein (CRP), Orosomucoid 1 and 2 (ORM1, ORM2), Prostaglandin E synthase (PTGES), and Signal transducer and activator of transcription 1 and 6 (STAT1, STAT6). While in females there was upregulation of Major histocompatibility complex, class II (HLA-DMB, HLA-DQB1), Signal transducer and activator of transcription 5B (STAT5B), and the nuclear factor kappa B subunits 1 and 2 (NFKB1, NFKB2) (Fig. 4). Interestingly, Superoxide dismutase (SOD3) that protects tissue by eliminating superoxide radicals was upregulated only in female tissue (3.5-fold change, FDR 0.024).

## DISCUSSION

The most significant finding from this study is that intact looking cartilage taken from ACL surgeries, which had only been exposed to an injurious environment, shows signs of cartilage remodeling in both sexes even in the absence of apparent tissue degeneration. These changes occur shortly after injury, as 80% of the samples were taken between 1.5- and 3.5-months post injury.

Given the limitations in obtaining young and healthy human cartilage to be utilized as control for research, investigators have used biopsies from non-load-bearing areas obtained at the time of reconstructive surgery or non-damaged regions in injured or OA joints, as this tissue has not been directly damaged and looks macroscopically normal^21–23^. However, the joint is an organ system, with shared synovial fluid and with high crosstalk between tissues^25, 40^. Therefore, even if some tissues are macroscopically intact, subcellular and metabolic changes have been shown to occur in intact appearing knee cartilage from OA joints^26^. Supporting this, in early onset hip osteoarthritis due to hip dysplasia, undamaged articular cartilage already displays OA-like intracellular changes (cytoskeleton remodeling, integrin upregulation) even though the ECM still appears intact^24^. Moreover, others have found that changes in the pericellular matrix (PCM) precede alterations in the ECM in a PTOA model^19^. These findings suggest that the impact triggered by injury originates at the cellular level slowly progressing to the PCM and likely taking an extended period to develop detectable signs at the ECM level. Therefore, to engineer efficient strategies to delay or even avoid PTOA, the early response after injury should be targeted. Despite the post-mortem time and biological heterogeneity of the normal samples, we were still able to detect signs of ECM remodeling in injured samples and some sex-specific responses. Of note, obtaining fresh and viable normal cartilage from young post-mortem donors requires a waiting time for passing serology screening.

Even though the injured tissue used in this study was macroscopically intact, we found evidence of ECM remodeling in both sexes. There was a decrease in levels of ACAN, COMP, MATN3, CILP, FN-1, and GDF10. On the other hand, there was an increase in COL1, 4, 5, 6, 14, the matrix metalloproteases MMP3, 14, 19, and ADAM10. The release of ACAN and COMP from the ECM is a hallmark of cartilage turnover and OA. COMP levels increase in serum and synovial fluid after knee injury^41, 42^ and in OA^42–44^ as a result of being released from the ECM, consistent with our findings of reduction in COMP levels in injured cartilage matrix. We observed an increase in COL1A1 and COL1A2 which is not a major hyaline cartilage component but rather more relevant to fibrocartilage as an attempt to repair tissue after injury^45^. Another protein involved in cartilage homeostasis, repair, and collagen accumulation is TGFβ1^46^, which we found upregulated in both sexes. Among many other effects, activation of TGFβ1 causes upregulation of fibronectin (FN-1) that can lead to fibrosis when FN-1 is produced in excess^47^. Contrastingly, we observed a decrease in FN-1 levels in both sexes, suggesting that the triggered response is not necessarily fibrotic. Interestingly, both sexes showed upregulation of DDR2 and CD44 in injured cartilage. DDR2 is a receptor tyrosine kinase that binds collagen triple helix and senses microenvironmental changes, such as collagen remodeling^48, 49^ and has been implicated in OA^49^. In fact, a mouse PTOA model with DDR2 haploinsufficiency shows a significant reduction in cartilage degeneration^50^. On the other hand, CD44 is the receptor for hyaluronic acid with essential roles in cell-matrix adhesion. Although it is expressed in normal cartilage, its expression can be a predictor of OA progression^51^.

In terms of sex-specific analysis, male cartilage showed a greater number of ECM-related proteins affected with injury compared to female cartilage. Most evident examples in injured male cartilage were the upregulation of the matrix proteases MMP2 and ADAM9, upregulation of several members of the serpin family which are serine protease inhibitors, and some are involved in chondrocyte differentiation^52, 53^ (SERPIN A1, A3, D1, G1). Furthermore, injured male cartilage showed downregulation of collagen IX (COL9A3) which is important for the structural arrangement of collagen II network and is downregulated in OA cartilage^21^, cartilage intermediate layer protein 2 (CILP2) that is downregulated after mechanically-induced joint destabilization^54^, and the tissue inhibitor of metalloproteinase 3 (TIMP3) which is a potent inhibitor of aggrecanases^42^.

Our analysis did not detect common cytokines involved in OA, such as IL-1β or TNF-α. Since we lack data on inflammatory serum markers from the individuals included in this study, we cannot conclude whether the acute inflammatory phase had already passed or whether there was no production of common pro-inflammatory cytokines locally in cartilage. However, there were several proteins involved in inflammatory and immune response in both male and female cartilage that were detected in our analysis. Examples of these were members of the complement system, such as the complement components 1, 4, 5, and 8. Members of the complement system are expressed by several cell types including articular chondrocytes^55^, and its upregulated expression has been detected in the synovial fluid of OA joints^56^. Injured male cartilage had 16 core components of the complement system upregulated and injured female tissue had 11. Moreover, only injured male cartilage showed upregulation of C-reactive protein (CRP), Prostaglandin E synthase (PTGES), Orosomucoid 1 and 2 (ORM1, ORM2), and the Signal transducer and activator of transcription 1 and 6 (STAT1, STAT6). On the other hand, only injured female cartilage showed upregulation of the Nuclear factor kappa B subunits 1 and 2 (NFkB1, NFkB2), which are downstream products of inflammatory cascades and early cartilage response to injury^57^.

The response to injury and repair involving collagen synthesis translates into high bioenergetic demands, which can be sourced from chondrocyte metabolism. Ample evidence shows that cellular metabolism is detrimentally affected in OA^58^. In fact, shifting from the TCA cycle to glycolysis has been associated with inflammation and OA progression^59^. Chondrocytes need to keep a steady ATP production to maintain the ECM homeostasis as collagen production depends on glucose consumption^60^. Therefore, it was interesting to find that several metabolic-associated proteins were upregulated in both sexes. Examples are Hexokinase (HK1), Citrate synthase (CS), Isocitrate dehydrogenase 2 (IDH2), and Oxoglutarate dehydrogenase (OGDH). These proteins have been observed to be upregulated in OA^58^.

Limitations of our study include the small sample size per sex, which reduces the ability to control for potential confounders like injury mechanisms and biological heterogeneity given by genetics and life style factors. However, we were still able to detect subtle but meaningful sex-specific responses. Despite the interconnectivity of energy consumption and ECM synthesis, our data suggests that male and female cartilage responses to injury diverge in their targeted pathways. We found that the molecular functions enriched in males were associated to ECM binding and structural constituents, while molecular functions in females were associated to energy use and protein synthesis, possibly prioritizing cellular resources for repair differently, or to prioritize maintenance.

Our data showed that chondrocyte metabolism in *injured* cartilage differed between males and females. The KEGG pathway analysis revealed that female cartilage had enrichment in *“Metabolic pathways,”* “*Fatty acid metabolism*,” “*Fatty acid degradation*,” and “*Citrate (TCA cycle)”* pathways. In addition, Serine hydroxymethyltransferase 2 (SHMT2), which catalyzes the mitochondrial conversion of serine to glycine, required for TGFβ-induced collagen synthesis, is upregulated in female injured cartilage only. Jain et al, in 2024, demonstrated that there are sex differences in energy metabolism in chondrocytes from OA knees^61^. While male chondrocytes prioritized glucose consumption and increased lactate production, female chondrocytes had a significantly higher fatty acid consumption, oxygen consumption rate, electron transport chain activity, and mitochondrial respiratory capacity, however, with no sex differences in number and area of mitochondria per cell^61^. Metabolic sex differences also have been found in human synovial fluid after knee injury, where levels of cervonyl carnitine were higher in females compared to males^62^. Carnitines are required to transport fatty acids into the mitochondria. These findings confirm our own findings of metabolic differences between males and females. The higher levels of estrogen present in women before menopause compared to men may contribute to promoting the expression of proteins related to fatty acid oxidation and metabolic pathways such as the adenosine monophosphate-activated protein kinase (AMPK) that can be activated by estrogen^63^. Indeed, all our samples were pre-menopausal aged.

Because glycolysis increases with inflammation, the lactate production from this metabolic pathway increases acidity. Females, but not males, showed upregulation of the Carbonic anhydrase 1 (CA1), which is crucial to maintaining a stable pH. Acidity can exacerbate the production of reactive oxygen species and therefore increase oxidative stress. In fact, reducing free radical production prevents chondrocyte death induced by impact injury^64^. It is interesting that only females had upregulation of CA1 and Superoxide dismutase 3 (SOD3), which is an enzyme that neutralizes superoxide radicals. This could be related to estrogen, as an increase in SOD activity in response to 17-β Estradiol has been reported^65^.

Another limitation of our study is that normal cartilage samples included condylar areas besides the intercondylar areas included in injured cartilage (see methods), which could affect the protein profile. However, our data showing upregulation of typical markers found in matrix remodeling due to inflammation (downregulation of ACAN and COMP, upregulation of matrix proteases, complement system and inflammatory markers) are not solely explained by differences in location.

Further investigation is needed to determine which of these molecular changes lead to cartilage degeneration and potentiate development of PTOA. In terms of sex-specific analysis, male and female cartilage share mostly similar responses to ACL injury, with similar changes at the ECM level and subtle sex-specific signatures. Nevertheless, within this small group of targets, we observed that females were enriched in proteins and pathways related to metabolism and protein synthesis, while males were enriched in proteins and pathways related to ECM and cell-ECM interaction. Additional studies are needed to define the impact of these sex differences over time; for example, whether they amplify at a later time point and lead to differential processes for cartilage degeneration that would need to be accounted for in the development of new strategies to mitigate PTOA risk after ACL injury.

## ACKNOWLEDGEMENTS

The authors thank Dr. Andrew Lemoff and Dr. Joseph Otto from the Proteomics Core Facility at UT Southwestern. This research was made possible through the support of AlloSource as they provided research consented human tissue. The authors also thank the donors’ families.

## AUTHOR CONTRIBUTION

Conception and design: PAH, SBS. Analysis and interpretation of the data: PAH, CYH, CX, CRC. Provision of study materials: SBS, CRC, JLP. Technical and logistic support: MVV. Obtaining of funding: PAH, SBS. Statistical expertise: CYH, CX. Drafting the article, critical revision, and final approval: all authors.

Generative AI and AI-assisted technologies were NOT used in the preparation of this work.

---

**Supplementary Table 1:** Demographics of the normal samples used.

| Sample Type | Sex | Age |
| --- | --- | --- |
| Normal | M | 37 |
| Normal | M | 38 |
| Normal | M | 39 |
| Normal | M | 39 |
| Normal | M | 39 |
| Normal | F | 37 |
| Normal | F | 38 |
| Normal | F | 38 |
| Normal | F | 39 |
| Normal | F | 39 |

